# Extreme rainfall drives early onset cyanobacterial bloom

**DOI:** 10.1101/570275

**Authors:** Megan L. Larsen, Helen M. Baulch, Sherry L. Schiff, Dana F. Simon, Sébastien Sauvé, Jason J. Venkiteswaran

## Abstract

1. The increasing prevalence of cyanobacteria-dominated harmful algal blooms is strongly associated with nutrient loading and changing climatic patterns. Changes to precipitation frequency and intensity, as predicted by current climate models, are likely to affect bloom development and composition through changes in nutrient fluxes and water column mixing. However, few studies have directly documented the effects of extreme precipitation events on cyanobacterial composition, biomass, and toxin production.
2. We tracked changes in a eutrophic reservoir following an extreme precipitation event, describing an atypically early toxin-producing cyanobacterial bloom, successional progression of the phytoplankton community, toxins, and geochemistry.
3. An increase in bioavailable phosphorus by more than 27-fold in surface waters preceded notable increases in *Aphanizomenon flos-aquae* throughout the reservoir approximately 2 weeks post flood and ~5 weeks before blooms typically occur. Anabaenopeptin-A and three microcystin congeners (microcystin-LR, -YR, and -RR) were detected at varying levels across sites during the bloom period, which lasted between 3 and 5 weeks.
4. *Synthesis and applications:* These findings suggest extreme rainfall can trigger early cyanobacterial bloom initiation, effectively elongating the bloom season period of potential toxicity. However, effects will vary depending on factors including the timing of rainfall and reservoir physical structure. In contrast to the effects of early season extreme rainfall, a mid-summer runoff event appeared to help mitigate the bloom in some areas of the reservoir by increasing flushing.

## Introduction

Harmful algal blooms (HABs) have increased in intensity, frequency, and distribution across Canada and the world (Ho, Michalak, & Pahlevan, 2019; Orihel et al., 2012; Pick, 2016). In Ontario alone, the number of reported HAB events between 1994 to 2009 increased by 22×, and, by 2009, nearly half of those reported were dominated by cyanobacteria (Winter et al., 2011). This increase in cyanobacterial growth and dominance has been strongly attributed to increased nitrogen (N) and phosphorus (P) availability (Paerl and Huisman 2009, O’Neil et al. 2012, Sukenik et al. 2015, Paerl et al. 2016) and more recently, climatic shifts. For example, thermal regime changes have resulted in earlier and warmer water temperatures (O’Reilly et al., 2015; Richardson et al., 2017), increasing the competitive advantage of certain bloom-forming cyanobacteria over other organisms (Jöhnk et al., 2008; Paerl & Huisman, 2009). Despite predicted climatic changes in both temperature and precipitation patterns (IPCC, 2007; McDermid, Fera, & Hogg, 2015), less attention has been given to the importance of precipitation regime changes, which may have an even greater impact on cyanobacteria bloom dynamics due to mixing and nutrient cycles (reviewed in Reichwaldt and Ghadouani 2012).

Extreme rainfall events, both in terms of frequency and intensity, are one of the many climatic components predicted to change over the next quarter century (Soulis, Sarhadi, Tinel, & Suthar, 2016). Depending on soil conditions, high intensity precipitation across a watershed may mobilize large quantities of sediment as well as soil-bound and soluble nutrients resulting in notable increases to in-lake nutrient bioavailability and suspended solids (Bouvy et al., 2003; Wood et al., 2017). Increased stream flow will also increase water column mixing, alter flushing rates and residence times, and weaken vertical stratification (Bouvy et al., 2003; Reichwaldt & Ghadouani, 2012) - all factors known to influence cyanobacteria bloom development (Jacobsen & Simonsen, 1993; Mitrovic, Oliver, Rees, Bowling, & Buckney, 2003; Wood et al., 2017). However, the degree to which those factors will affect primary production in a reservoir is directly related to event timing (e.g. early vs. late summer) and the zonal gradient created by basin morphometry differences (Kimmel & Groeger, 1984).

Reservoirs naturally exhibit spatial heterogeneity in physical, chemical, and biological properties (Thornton, Steel, & Rast, 1996) as the narrow, channelized flow from the catchment (riverine zone) transitions (transitional zone) to a broader and deeper, lake-like system (lacustrine zone). Primary production along the riverine-transitional-lacustrine gradient is primarily mediated by light and nutrient availability, among other interacting factors (Kimmel & Groeger, 1984). However, large precipitation events have the potential to disrupt characteristic primary production patterns in these zones, especially if the event introduces sizable nutrient loads, increases flushing rates, or affects other water-column properties prior to or during a bloom period. For example, Wood et al. (2017) reported decreased cyanobacterial biomass following an extreme rainfall event that led to water column cooling and destratification in a shallow, eutrophic New Zealand lake. Varying responses to extreme rainfall should be anticipated given the importance of site and event-specific factors.

Changing cyanobacterial bloom dynamics and biomass are a great concern to water managers. Several bloom-forming cyanobacterial species such as *Aphanizomenon flos-aquae*, *Microcystis aeriginosa*, and *Anabaena (Dolicospermum) flos-aquae* synthesize an array of bioactive compounds that pose acute, chronic, and potentially fatal health risks to humans and other organisms through dermal contact, inhalation, and/or ingestion of contaminated waters (Chorus & Bartram, 1999; Chorus, Falconer, Salas, & Bartram, 2000; Codd, 2000). Metabolite production as well as the composition of toxins released are complex and tied not only to the cyanobacterial species, successional patterns, and biomass, but also to various environmental parameters including N, P, temperature, light, pH, salinity, and micronutrients (Chorus & Bartram, 1999), which may all be altered by intense rainfall events (Reichwaldt & Ghadouani, 2012). To date, only limited information exists regarding how toxicity may change as a result of precipitation events.

Here, we document the effects of an extreme early summer rainfall event on cyanobacterial bloom and toxin dynamics in a eutrophic reservoir and contrast these effects to a second, smaller mid-summer rainfall event. Conestogo Lake (Ontario, Canada; Fig. 1), has experienced recurring, late-summer cyanobacterial blooms dominated by *Aphanizomenon flos-aquae* since 2004. On 23^rd^ June 2017, the upper Conestogo watershed experienced a record rainfall event (daily total 78 mm measured at Conestogo Dam; https://www.grandriver.ca/en/our-watershed/Record-Rainfall-Flood-June-2017.aspx) that resulted in severe flooding and replaced ~80% of the water in Conestogo Lake during this 2-day event (Grand River Conservation Authority (GRCA) 2018). We first evaluate how this extreme event affected the reservoir by characterizing the physical and chemical changes across spatial zones. We then follow the trajectory of bloom development and toxicity through the season, to better understand how hydrological events can alter bloom risk.

**Fig. 1.**
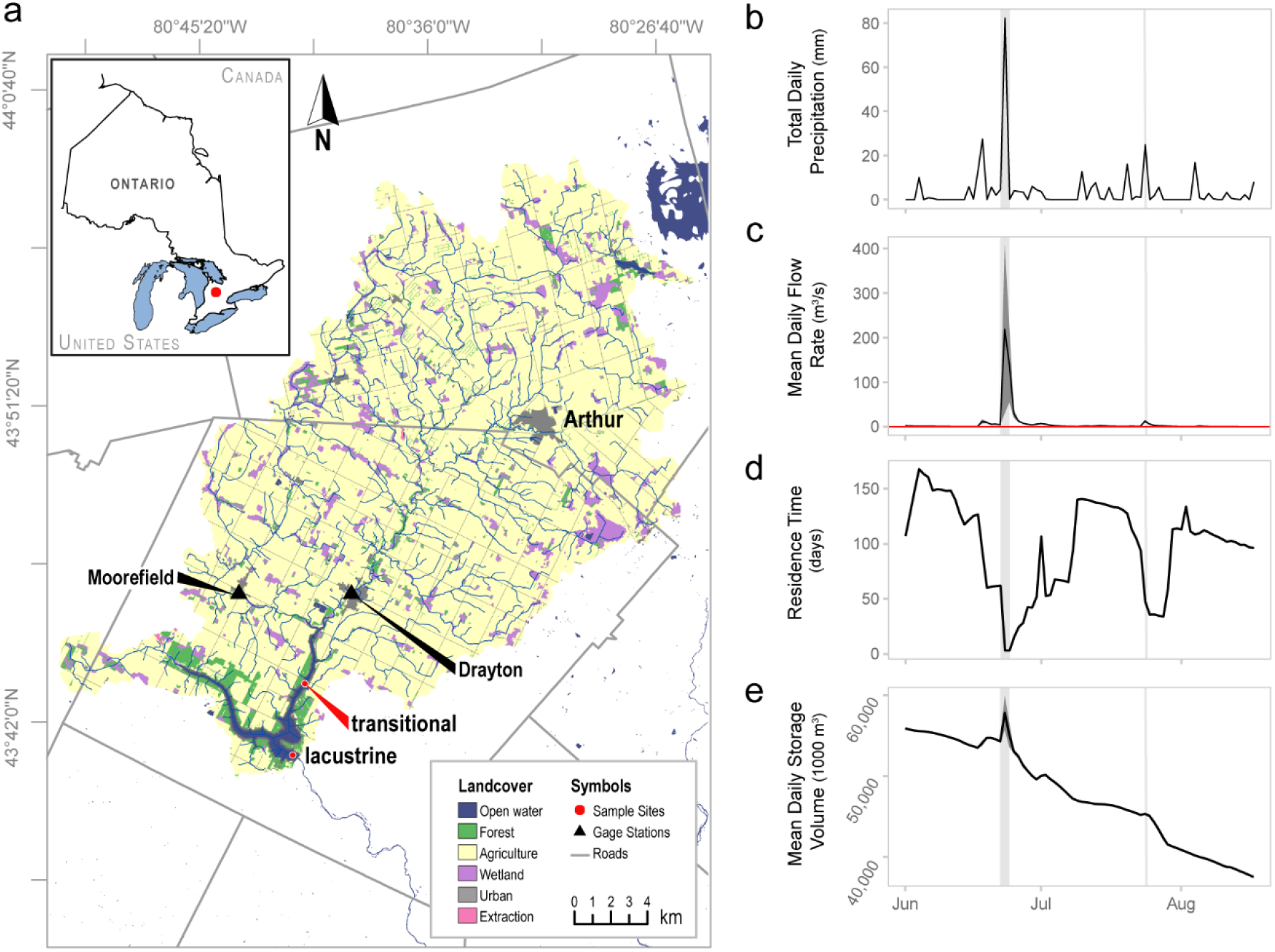
(*a*) Conestogo Lake (Mapleton Township, Ontario, Canada) and its associated watershed. Sampling began on 21 Jun 2017 (DOY 172) in the lacustrine zone and on 05 Jul (DOY 186) in the transitional zone. Total daily precipitation (mm) measured in upper Conestogo watershed (*b*, gauge in Arthur, ON) was directly related to the Conestogo River mean daily flow rate (*c, n* = 48) into the reservoir (gauges in Drayton and Moorefield, ON) and the calculated reservoir residence time (*d*). The average flow rate during the drawdown period (May – September; red line, *c*) is typically near 0.01 m^3^ s^-1^. The extreme precipitation event on 23-24 Jun (DOY 173-174; grey vertical bar) directly corresponds to in increased mean daily flow into the Conestogo Reservoir, increased reservoir storage, and reduced residence time. Dark gray area around the mean represents the standard deviations of daily measurements (flow, *n* = 48; storage, *n* = 24). A secondary rainfall event (~25 mm) on 24 Jul (DOY 205) also impacted flow rates into the reservoir. This figure contains information made available under GRCA’s Open Data Licence v3.0 (*a*) and v2.0 (*b* - *e*).

## Materials and methods

### Sample collection

We sampled a lacustrine (43.676518, −80.718752; max depth = 17 m) and transitional (43.708102, −80.712224; max depth = 9 m) site in Conestogo Lake once per week from 21 Jun to 17 Aug 2017. Water column profiles and samples for chemical analysis were collected at the lacustrine site on 21 Jun (DOY 172) with biological sampling beginning on 05 Jul (DOY 186). Sampling in the transitional zone began on 05 Jul, 2 weeks after the extreme rainfall event. (Figure 1*a*). We measured water column profiles at 0.5 m increments using an EXO2 sonde (YSI, Yellow Springs, OH) equipped with chlorophyll-a (μg L^-1^), pH, temperature (°C), and dissolved oxygen (DO, mg L^-1^) sensors. Photosynthetically active radiation (PAR) in the water column was also measured in 0.5 m increments using a LI-COR^®^ Underwater Quantum Sensor (LI-250A, LI-192, Lincoln, NE USA).

For all chemical analyses, discrete water samples were collected at 2 m and 7 m below (lacustrine only) the water surface, and 0.5 m from the bottom using a Van Dorn bottle. Unfiltered whole water samples at each depth were portioned into acid-washed 50 mL centrifuge tubes for total phosphorus (TP, μg L^-1^) and into a 125 mL dark bottle for phytoplankton enumeration (2 m only). Additional aliquots were field-filtered using a 0.45 μm syringe filter (Whatman) for soluble reactive phosphorus (SRP, μg L^-1^), total dissolved phosphorus (TDP, μg L^-1^), ammonium (NH_4_^+^, mg N L^-1^), total dissolved nitrogen (TDN, mg N L^-1^), anions (NO_2_^-^, NO_3_^-^, mg N L^-1^; Cl^-^, SO_4_^-^, mg L^-1^), total dissolved iron (TDFe), mg L^-1^), and cations (Mg^2+^, Ca^2+^, mg L^-1^). All samples were transported on ice and chemical samples stored frozen until analysis at the University of Waterloo (Waterloo, ON) or Centre for Cold Regions and Water Science (Waterloo, ON).

Samples for total (*i.e*., intracellular and extracellular) metabolites were pooled from four discrete depths across the site-specific photic zone (2× Secchi depth). The method serves to sample the photic zone as a mechanism of estimating all possible toxins in the surface and near-surface. The estimated photic zone was split into four equal segments, sampled at each segmented depth using a Van Dorn bottle, then pooled in a bleach-sterilized bucket. For example, a Secchi depth measured at 1 m would be multiplied by two then divided by four to create four discrete samplings at 0.5, 1, 1.5, and 2 m. Each pooled sample was collected in an amber Nalgene™ polyethylene terephthalate glycol (PETG) bottle to limit adsorption (Fisher reference: 322021-0125). All samples were stored frozen at −20 °C until analysis at the Université de Montréal (Montréal, QC).

### Chemical analyses

SRP, TDP, and TP concentrations were measured on a Cary 100 UV-Vis spectrophotometer (Santa Clara, CA) following off-line persulfate digestion (TP samples only) as per standard methods (Eaton, Clesceri, Greenberg, & Franson, 1998). Each N parameter was analyzed on a separate instrument with NH_4_^+^ and TDN analyzed on a Westco Smartchem Analyzer (Unity Scientific, Milford, MA) and Shimadzu TN-L analyzer (Kyoto, Japan), respectively. Anion concentrations including NO_2_^-^ and NO_3_^-^ were measured on a Dionex Ion Chromatogram ICS-200 (Waltham, MA) while cations and TDFe were measured on a Perkin Elmer Optima 8000 ICP-OES (Waltham, MA).

### Phytoplankton identification and enumeration

Phytoplankton enumeration was completed by D. Findlay (Plankton-R-Us, Winnipeg, MB, http://www.plankton-r-us.ca) as per Findlay & Kling (2003). Each sample was preserved with 4% Lugol’s iodine, gravity concentrated 5-fold after 24 hrs, and finally, stored at 4 °C for ~2 months until analysis. Enumeration was completed on an inverted microscope at 125x, 400x, and 1200x with phase contrast illumination using a modified Ütermohl technique (Nauwerk, 1963) on 10 mL aliquots of preserved sample. Only cells with viable chloroplasts were enumerated.

Cell counts were converted to mg wet weight biomass (mg m^-3^) by approximating cell volume, which were obtained by measurements of up to 50 cells of an individual species and applying the geometric formula best fitted to the shape of the cell (Vollenweider, 1968). A specific gravity of 1 was assumed for cellular mass (D. Findlay, pers. communication).

### Cyanobacterial metabolite analysis

Samples were prepared and screened for each of seventeen cyanobacterial compounds (Table S1) as per Fayad *et al*. (2015) via on-line solid phase extraction ultra-high performance liquid chromatography high resolution mass spectrometry (SPE-UHPLC–HRMS) using standards purchased from Enzo Life Science, Abraxis, or Cyano Biotech GmbH. The limit of detection (LOD) and limit of quantification (LOQ) were calculated for every batch of samples. Only results that exceeded the LOQ were included in our analysis. Therefore, some samples may have had detectable, but unquantifiable toxin concentrations.

### Environmental and dam-related data

Meteorological and other reservoir-related data for this study were obtained from the GRCA’s online data portal (https://data.grandriver.ca/). Because GRCA data are provisional, all data were passed through quality assurance and quality checking metrics to ensure sound data quality before proceeding with analysis. Spatial data were obtained from open source portals at the GRCA, the United States Geological Society (https://www.usgs.gov/), Statistics Canada (https://www12.statcan.gc.ca/), and the Ontario Ministry of Natural Resources (land-use; https://www.ontario.ca).

### Data analysis and statistics

All data analysis was completed in R version 3.6.1 (R Core Team, 2018). Sonde-derived water column profiles as well as the discrete chemical profiles were constructed using a multilevel B-spline interpolation from the *MBA* package (Finley, Banerjee, & Hjelle, 2017) and were adjusted to reflect the reservoir stage elevation at the time of sampling.

To compare phytoplankton community composition with environmental variables, we used Principal Components Analysis (PCA) and redundancy analysis (RDA) including environmental variables from the epilimnion and 0.5m above the sediment-water interface. Species biomass data were Hellinger transformed as recommended for species linear ordinations (Legendre & Gallagher, 2001), which normalizes for both the site and species biomass needed for RDA. Only species that appear more than two times in the dataset were included in these analyses. The environmental matrix included all chemical parameters at 2 m depth (epi) and those from 0.5 m above the sediment-water interface (sed), sonde-derived measurements for the epilimnion and hypolimnion (temperature, pH, DO) as well as physical lake parameters (thermocline depth and metalimnion layer depth; calculated using the *rLakeAnalyzer* package (Winslow et al., 2019)). We constructed PCA biplots with either species vectors which exceeded a cumulative goodness-of-fit of at least 0.6 in the ordination plane or significant environmental parameters. The higher the goodness-of-fit, the better the species is fitted on the corresponding axis. To test for the significance of environmental factors affecting the phytoplankton community, permutation tests (permutations = 999) on the models were completed using *anova.cca* from the package *vegan*. Epilimnetic chemistry, biomass, and toxin trends were analyzed using repeated measures ANOVA (RM-ANOVA) with a generalized least squares model (*gls*) from the *nlme* package (Pinheiro, Bates, DebRoy, Sarkar, & R Core Team, 2018)where time and site were fixed effects and the correlation matrix was selected based on Akaike Information Criterion (AIC). To account for temporal autocorrelation, we nested time within site. Finally, we tested for the correlation of biomass with toxin concentrations using Kendall’s rho.

## Results

### Extreme rainfall disturbance in early summer

Record rainfall (daily range 40 - 120 mm) was recorded across the Upper Conestogo watershed on 23 Jun 2017 (DOY 174) and led to widespread flooding. This event was the largest single rainfall in the Grand River Watershed (Luther Marsh station; ~50 km distant; records since 1950; https://www.grandriver.ca/en/our-watershed/Record-Rainfall-Flood-June-2017.aspx).

Mean daily flow rates into Conestogo Lake via the Conestogo River increased from 4.56 m^3^ s^-1^ to 219 m^3^ s^-1^ (max 521 m^3^ s^-1^) in the two day period, which consequently increased the total reservoir volume by 3.5 Mm^3^,and is estimated to have replaced ~80% of the total reservoir volume (Fig 1*b-e*) (GRCA 2018). Within several days, the reservoir storage and drawdown returned to levels targeted for regular reservoir operation.

Reservoir volume replacement restructured the physical water column. Both the transitional and lacustrine zones were well mixed, isothermal, and oxygenated immediately following the event (Fig 2). Thermal stratification did not return to either location during the sampling period. Intermittent hypolimnetic hypoxia (DO < 2 mg L^-1^) developed in the transitional zone while sustained hypoxia developed in the lacustrine within two weeks of the flood event (05 July, DOY 186). Lacustrine SRP, TDP, and TP increased between 2-33× across the sampled depths (2m: 7 to 27×, 7m: 9 to 33×, bottom [16m]: 2 to 3×; Figs. 2, 3) before and after the extreme rainfall. Nitrogen concentrations at all measured depths were only marginally different post-rainfall and maintained concentrations near 5.5 mg N L^-1^ for NO_3_^-^ (mean prior = 4.89 mg N L^-1^; mean after = 5.45 mg L^-1^) and TDN (mean prior = 5.63 mg N L^-1^; mean after = 5.71 mg L^-1^) (Fig. 2).

**Fig. 2.**
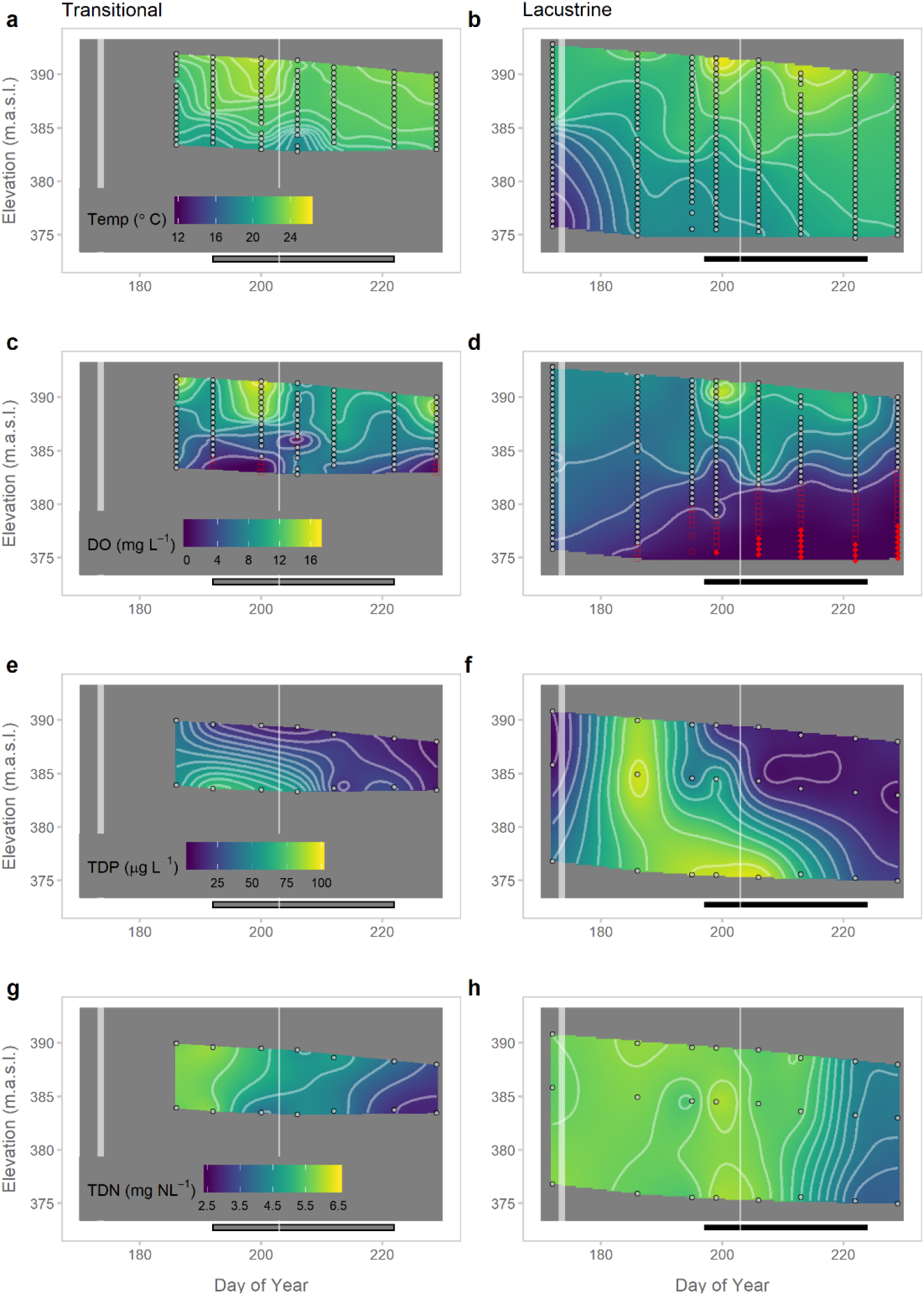
Temperature (*a-b*, °C) and dissolved oxygen (*c-d*, mg L^-1^) water column profiles collected at 0.5 m depth increments (grey points) in the transitional (*a, c*) and lacustrine (*b, d*) zones from 21 Jun 2017 (lacustrine only; DOY 172) to 17 Aug (DOY 229). Hypoxic (red squares) zones were detected at all sites, but anoxia (red triangles) was only detected in the lacustrine zone. TDP (*e-f*) and TDN (*g*-*h*) profiles collected at 2 m and 7 m below (lacustrine only) the surface and 0.5 m from the bottom. Profile depths at each site were standardized to the dam stage elevation (meters above sea level, m.a.s.l.) at the time of sampling. The grey regions in the x-y plane illustrate the interaction between time and reservoir depth at each site, where the x-axis zone indicates a later sampling date in the transitional and the y-axis grey zones illustrate differences in site depths with reference to the dam (max elevation = 393 m.a.s.l, dam outflow = 373.5 m.a.s.l.). The first white vertical bar corresponds to the record rainfall event on (23-24 Jun 2017; DOY 174-175) and the second to a 25 mm rainfall event on 24 Jul (DOY 205). The black horizontal bar corresponds to the bloom period in each of the two zones.

**Fig. 3.**
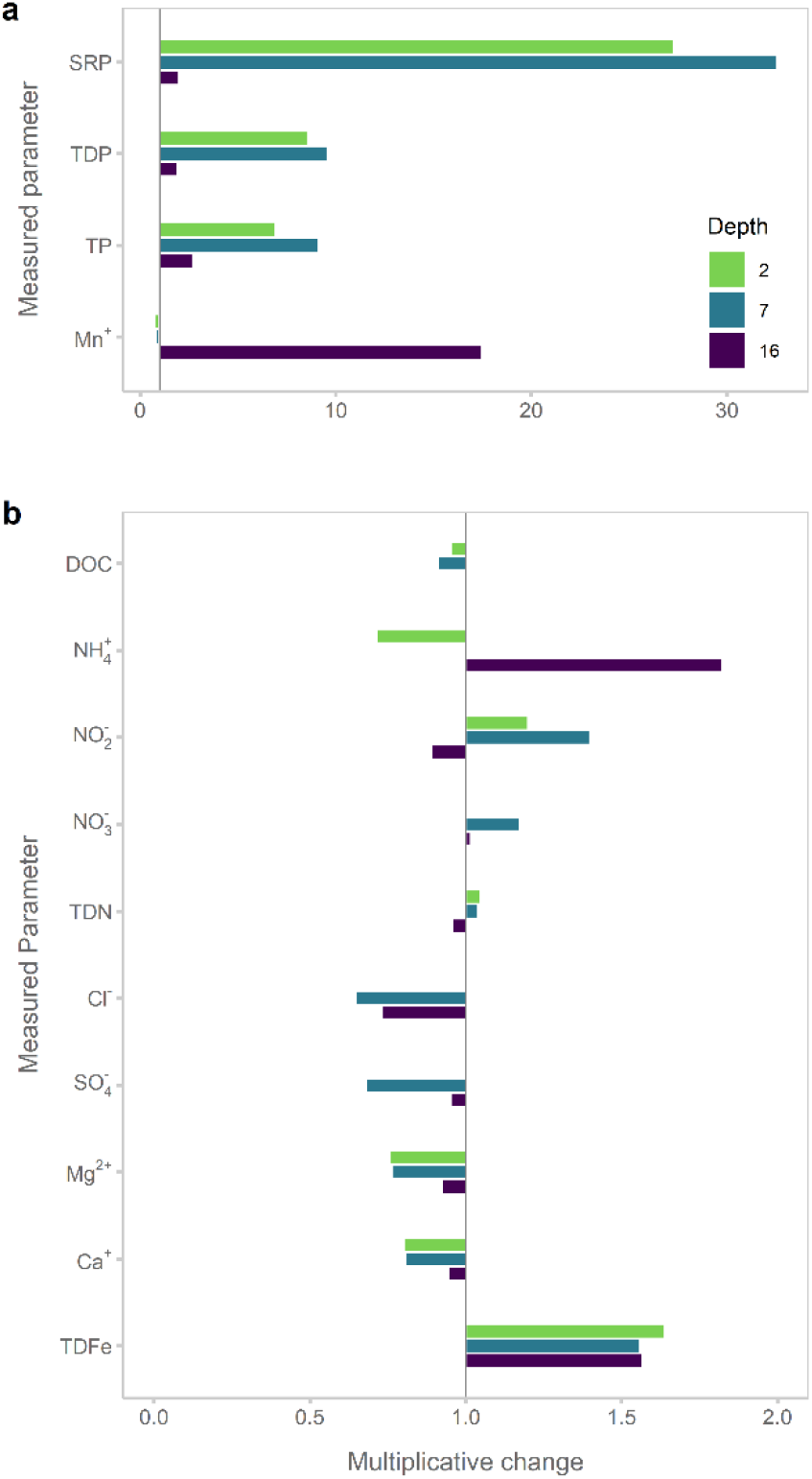
Multiplicative changes in phosphorus species and Mn (*a*), nitrogen species and DOC (*b*), anions (*c*), cations and TDFe (*d*) prior to (21 Jun 2017, DOY 172) and after the early summer flood event (05 Jul 2017, DOY 186) in the lacustrine zone. Samples were collected at 2 m depth (purple), 7 m depth (blue), and 0.5 m from sediment-water interface (16m; green) in the lacustrine zone. Values above 1 × (black horizontal line) correspond to increased concentrations while those below 1 × indicate reduced concentrations. Samples at 2m for anions (*b*[NO_3_^-^, Cl^-^, SO4^-^) were not analyzed.

### Dynamics following early summer extreme rainfall event

Mean surface water temperatures (0 - 2 m) across the reservoir ranged from 21.2 to 25.5 °C. Nutrient concentrations (both N and P) were, on average, higher in the transitional zone than the lacustrine zone and significantly changed through time (Table 1; RM-ANOVA). Epilimnetic chlorophyll a, dissolved oxygen (DO), and temperature did not significantly change through time across sites. As predicted, the transitional zone had higher concentrations of suspended solids in the water column and lower light availability as compared to the lacustrine immediately following the reservoir disturbance (Fig. 4).

**Fig. 4.**
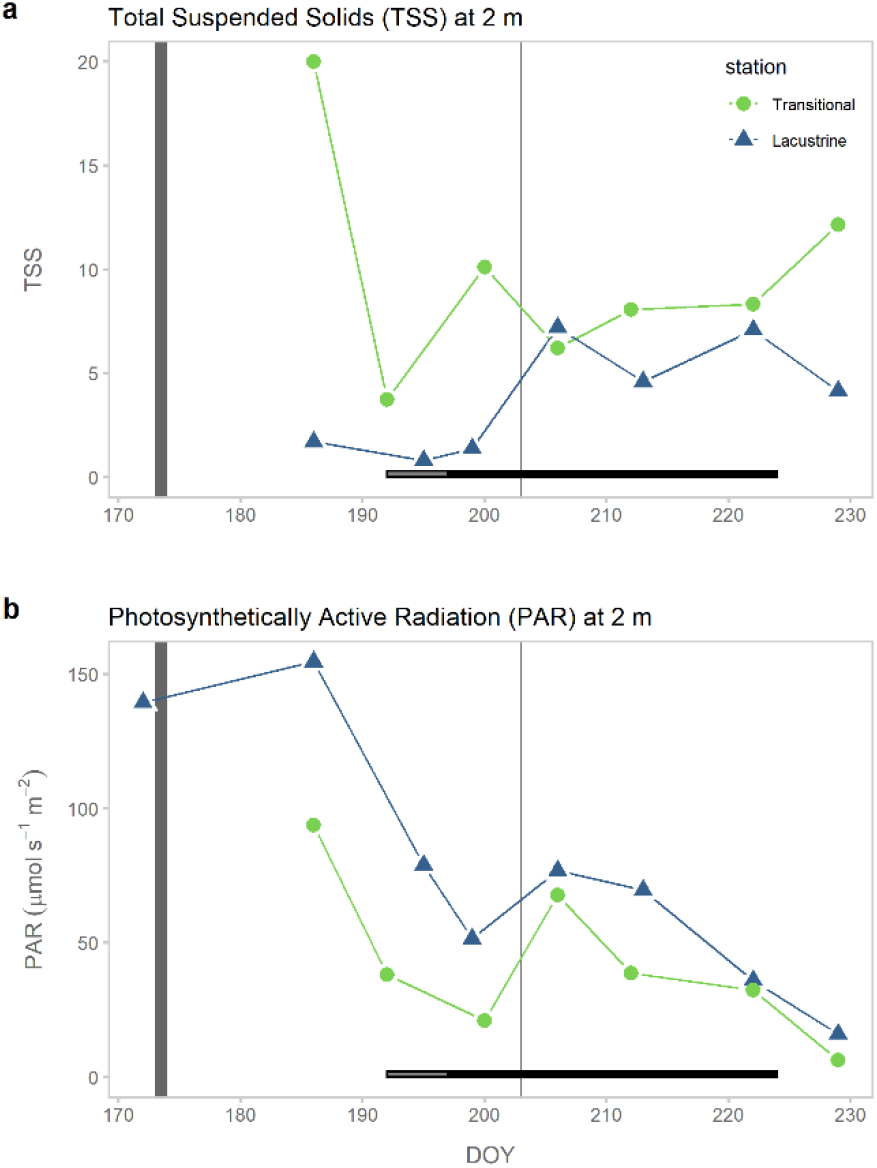
TSS (*a*) and light availability (*b;* photosynthetically available radiation (PAR) at 2 m in the transitional (green) and lacustrine (purple) zones. The horizontal bar corresponds to the cyanobacteria bloom period in the transitional (grey) and lacustrine (black) zones while the vertical bars are associated with the extreme rainfall event on 23 June (DOY 174) and the late summer event on 24 Jul (DOY 205).

**Table 1.** Post-flood mean (minimum - maximum) values for measured epilimnetic (2 m) characteristics in the transitional and lacustrine zones with RM-ANOVA *p*-values. RM-ANOVA tables are available in the supplement. Bold values are significant (*p* < 0.05).

|  | **Chl-a**<br>mg L<sup>-1</sup> | **DO**<br>mg L<sup>-1</sup> | **Temp**<br>°C | **NO<sub>3</sub>-N**<br>mg L<sup>-1</sup> | **TDN**<br>mg L<sup>-1</sup> | **SRP**<br>µg L<sup>-1</sup> | **TDP**<br>µg L<sup>-1</sup> | **PAR**<br>µmol s<sup>-1</sup> m<sup>-2</sup> |
| --- | --- | --- | --- | --- | --- | --- | --- | --- |
| **Transitional** | 6.8<br>(1.6 - 14.0) | 10.6<br>(7.9 - 14.5) | 22.8<br>(21.7 - 24.0) | 4.82<br>(3.99 - 5.65) | 5.32<br>(4.24 - 5.86) | 13.66<br>(1.52 - 65.2) | 28.76<br>(11.62 - 85.77) | 42.6<br>(6.33 - 93.9) |
| **Lacustrine** | 5.4<br>(2.2 - 10.1) | 8.6<br>(5.6 - 11.0) | 22.3<br>(21.2 - 23.2) | 4.37<br>(3.34 - 5.63) | 4.74<br>(3.39 - 5.85) | 5.89<br>(1.56 - 24.61) | 18.53<br>(7.79 - 41.59) | 69.1<br>(16.0 - 155) |
| **Intercept** | **0.001** | **< 0.001** | **< 0.001** | **< 0.001** | **< 0.001** | **0.01** | **0.001** | **< 0.001** |
| **time** | 0.415 | 0.882 | 0.234 | **< 0.001** | **< 0.001** | **0.014** | **0.007** | **0.003** |
| **site** | 0.586 | 0.204 | 0.244 | **0.001** | **0.004** | 0.196 | 0.206 | 0.064 |

Abbreviations: chlorophyll a (Chl-a), dissolved oxygen (DO), water temperature (temp), nitrate (NO<sub>3</sub>), total dissolved nitrogen (TDN), soluble reactive phosphorus (SRP), total dissolved phosphorus (TDP), photosynthetically reactive radiation (PAR)

#### Phytoplankton community composition

The transitional zone phytoplankton community was composed predominately of diatoms including *Stephanodiscus niagarae* (P5530) and *Cyclotella psuedostelligera* (P5508) and the dinoflagellate *Ceratium hirundenella* (P7644) with -3 approximately 5x higher total phytoplankton biomass in the transitional (total = 5742 mg m^-3^ wet weight) than the lacustrine (total = 1113 mg m^-3^ wet weight; 2 weeks post-extreme event, Fig. 6a). In contrast to the transitional zone, the lacustrine zone contained high relative abundances of Cryptophytes including *Cryptomonas rostratiformis* (P6565), *Cryptomonas erosa* (P6558), and *Rhodomonas minuta* (P6554) and Chrysophytes (*Ochromonas globose* [P4398]), all of which returned following the cyanobacterial bloom. Communities following the extreme rainfall event(DOY 186) were strongly associated with PAR and epilimnetic SRP, TDP, and TP. Communities in both zones gave way to cyanobacterial dominance (> 50% of community by biomass, Figs.S1 - S2) 20 - 24 days following the flood with apparent changes to epilimnetic temperature and pH associated with temporal community shifts and increased presence of the filamentous nitrogen-fixer, *Aphanizomenon flos-aquae* (Fig 5a; P1041).

**Fig. 5.**
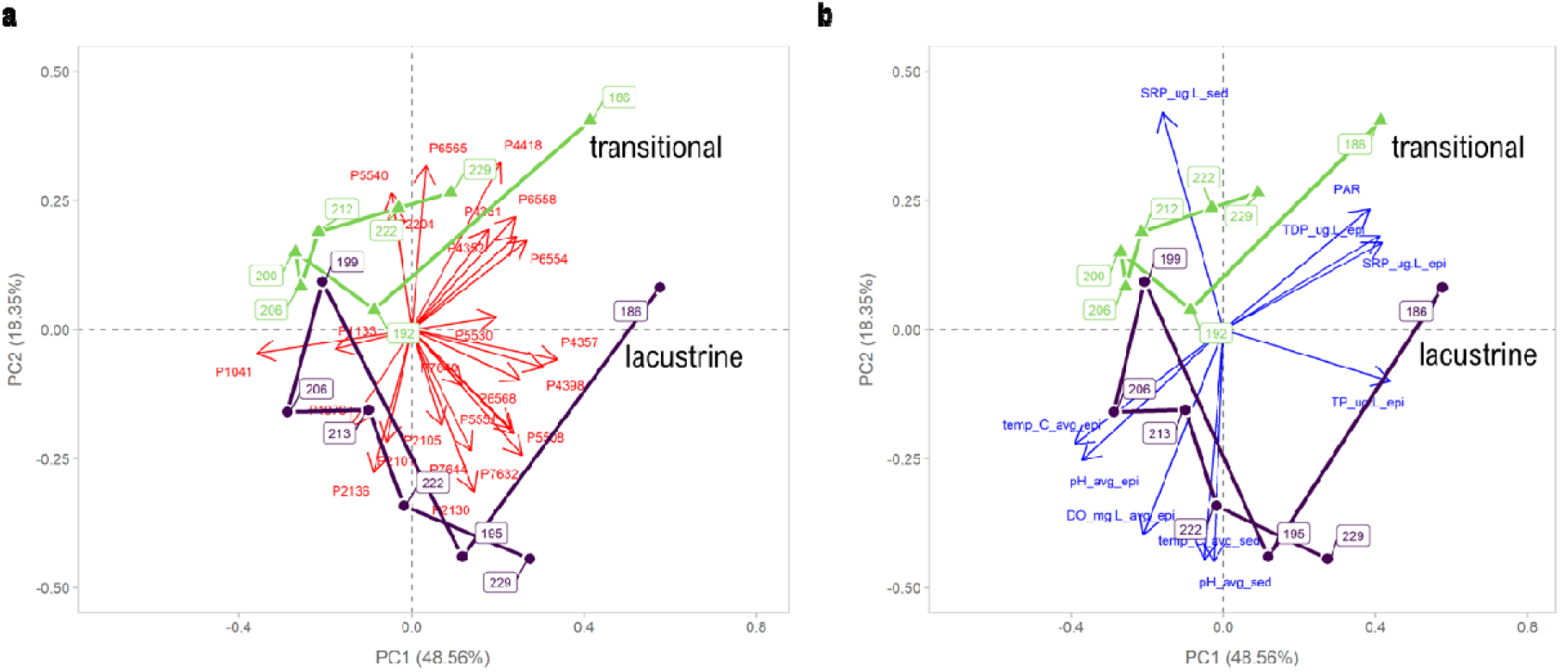
Principal component analysis (PCA) (scaling 1) of phytoplankton community, with the most influential phytoplankton species (*a*, Goodness of fit > 0.6) and significant (P < 0.05) and weekly significant (P < 0.1) environmental variables (*b*). The length of the vector is proportional to the importance of the descriptor to the sites. The day of year for each community are in boxes and are linked through time for each reservoir zone (lacustrine [purple], transitional [green]. Phytoplankton labels: P1041; *Aphanizomenon flos-aquae*, P2235; *Ankistrodesmus spiralis*, P6554; *Rhodomonas minuta*, P6558; *Cryptomonas erosa*, P4398; *Ochromonas globose*, P5515; *Fragilaria crotonensis*, P7644 *Ceratium hirundenella*. Environmental labels: SRP_ug.L_epi; epilimnetic soluble reactive phosphorus, SRP_ug.L_epi; sedimentary soluble reactive phosphorus, TDP_ug.L_epi; epilimnetic total dissolved phosphorus, TP_ug.L_epi; epilimnetic total phosphorus, PAR; photosynthetically active radiation, DO_mg.L_avg_epi; epilimnetic dissolved oxygen, pH_avg_epi; epilimnetic pH, pH_avg_sed; sedimentary pH, temp_C_avg_epi; epilimnetic temperature, temp_C_avg_sed; sedimentary temperature.

**Fig. 6.**
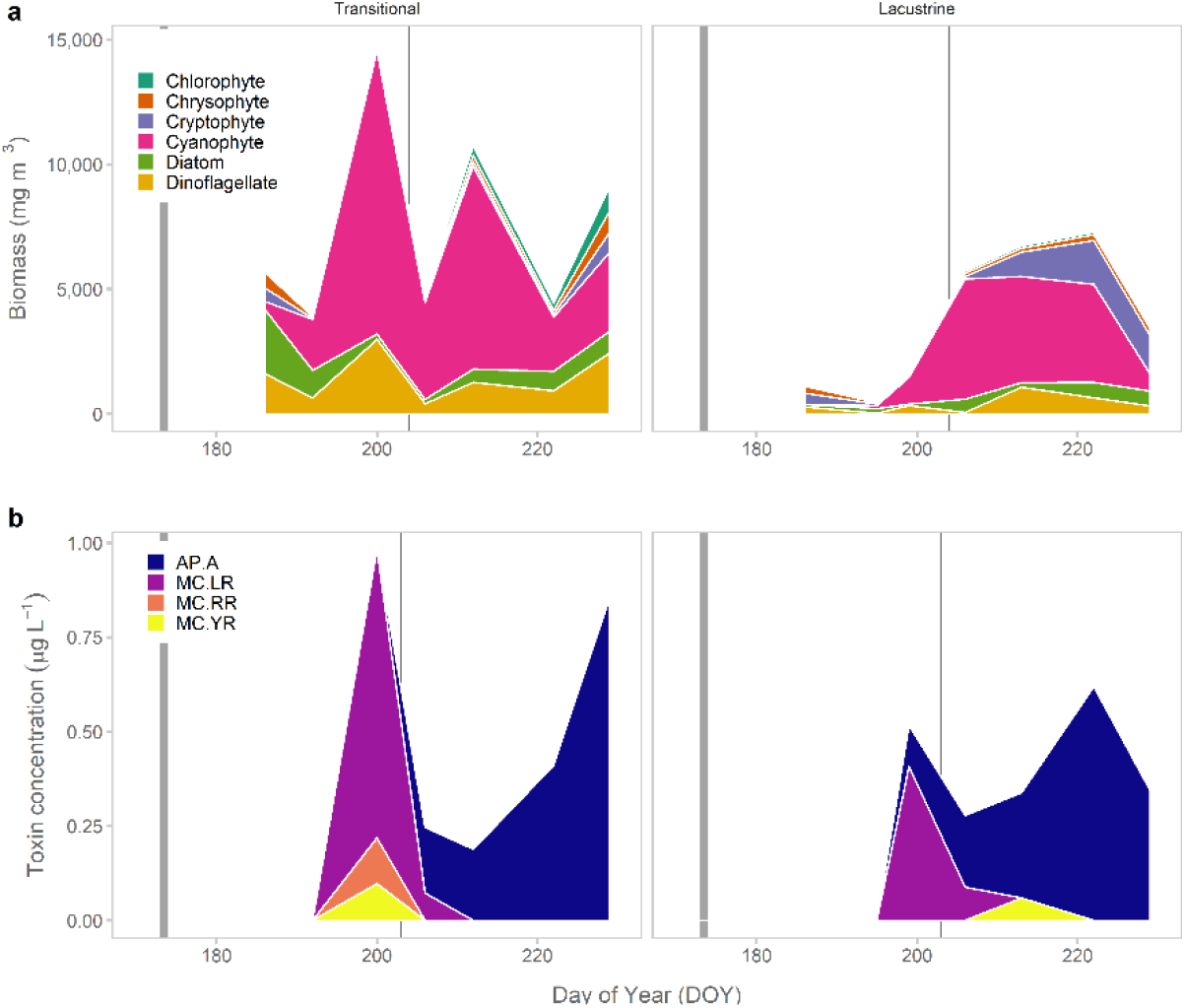
Phytoplankton biomass (*a*) and cyanobacterial metabolite concentration trends (*b*; exceeding the limit of quantification) for anabaenopeptin-A (AP.A) and microcystin-LR (MC.LR), -YR (MC.YR), and -RR (MC.RR) in the transitional and lacustrine zones. The flood event on 23-24 Jun (DOY 174-175 and secondary event on 25 July (205) are noted by vertical bars. *Aphanizomenon flos-aquae* dominated (> 50%) phytoplankton biomass in the transitional zone near 10 Jul 2017 (DOY 191) and in the lacustrine near 16 Jul 2017 (DOY 197). Only cyanobacterial species that were present in more than 2 samples are plotted above. Of the seven cyanobacterial species detected, various strains of *A. flos-aquae*, *A. flos-aquae (Dolichospermum flos-aquae*), *M. aeruginosa*, and *W. compacta* have been documented as potential toxin producers. Typical bloom onset occurs between DOY 220 – DOY 255 (S. Cooke, pers. communication).

#### Cyanobacterial dynamics

Cyanobacterial bloom duration, measured as the period during which cyanobacterial biomass comprised > 50% of the fractional biomass, persisted between 26 and 34 days and occurred 4 to 6 weeks earlier than has been previously documented by the GRCA (S. Cooke, pers. communication). Bloom initiation was 6 days earlier in the transitional zone as compared to the lacustrine. Mean total cyanobacterial biomass over the sampling campaign was 1550 ± 2920 mg m^-3^ and 591 ± 1290 mg m^-3^ in the transitional and lacustrine zones, respectively and was significantly driven by an interaction between SRP and surface temperatures (RM-ANOVA, F_*1, 6*_ = 6.84, *p* = 0.0399). There was no relationship between total cyanobacterial biomass and time or site (RM-ANOVA, *F_1,10_* = 0.086, *p* = 0.77). *Aphanizomenon flos-aquae* was the dominant species and represented 95-97% (by biomass) of the cyanobacterial fraction across all sites for the entirety of the sampling campaign. The highest measured *A. flos-aquae* biomass occurred on 19 July (DOY 200, 10,900 mg m^-3^, 4,500 cells mL^-1^) in the transitional and near 25 Jul (DOY 206; 4,710 mg m^-3^) in the lacustrine. *Woronichinia compacta*, *Anabaena (Dolichospermum) flos-aquae*, and *Microcystis aeruginosa* were minor contributors (< 5 %) to cyanobacterial biomass while *Anabaena* (*Dolichospermum) crassa, Planktolyngbya limnetica*, and *Pseudoanabaena sp*. were only observed once during the campaign at low biomass (< 8 mg m^-3^). GRCA managers have visibly recorded cyanobacterial blooms dominated by an *Aphanizomenon sp*. in Conestogo Lake from late August through September (DOY ~220 - 250) since 2004 (S. Cooke, pers. communication).

#### Cyanotoxin dynamics

Four of the 17 screened cyanobacterial metabolites were detected in Conestogo Lake. Detectable anabaenopeptin-A (AP-A), microcystin-LR (MC), -YR, and -RR were present in 87 % of the samples (*n* = 13) but were only quantifiable in 73 % (*n* = 11; Table 2). Total microcystin concentration did not exceed 1.0 μg L^-1^ and was comparable to previously measured concentrations during the summer months (Yakobowski, 2008). Metabolite type and concentration varied through time (RM-ANOVA, *F_3,44_* = 9.21, *p* = 0.001) and were different between the two sites (RM-ANOVA, *F_1,44_* = 6.55, *p* = 0.014). Quantifiable concentrations of all three microcystins were present in the transitional zone, while MC-LR and MC-YR were present in the lacustrine. MC-LR was the dominant microcystin variant across all sites with the highest quantified values in the transitional zone at peak *A. flos-aquae* and *M. aeruginosa* biomass (Fig 6*b*, Table 2). Further, MC-YR in the lacustrine did not correspond with the MC-LR peak. AP-A was present in both zones throughout the bloom period MC-LR concentration was significantly correlated with the biomass of *M. aeruginosa* (Kendall rho = 0.76, *p* = 0.03). No other toxin was correlated with the biomass of any other observed cyanobacterial species.

**Table 2.**
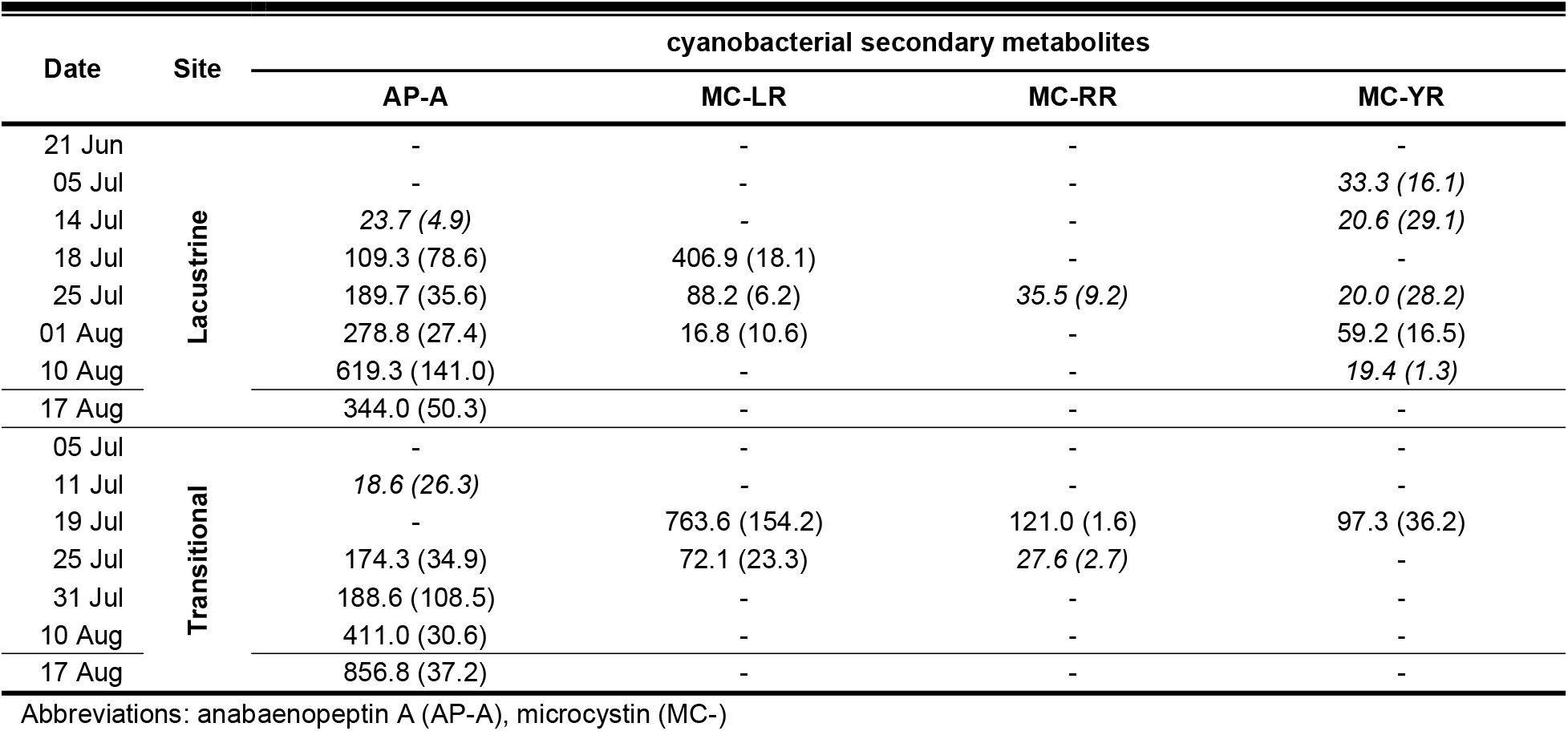
Quantified cyanobacterial metabolites across sampling sites as measured by LC-HMRS with values which exceeded the limit of detection (LOQ) presented as mean ng L^-1^ (standard deviation ng L^-1^), those in italics exceeded the limit of detection (LOD) but not the limit of quantification (LOQ), and those with (-) < LOD. The metabolites MC-LA, -LY, -LW, -LF, - HiIR, -HtyR, anatoxin, homoanatoxin, cylindrospermopsin, and cyanopeptolin A were not detected in samples.

#### Dynamics following smaller mid-summer runoff event

A second, much smaller and transient increase in flow to the reservoir occurred on 24 Jul (DOY 205), associated with an ~25 mm rainfall event occurring during already wet conditions (Fig. 1). Effects of this event were much more muted. Overall, there was a slight decrease in surface water temperatures and decrease in the extent of oxygen supersaturation in both reservoir zones,with minimal effects on nutrients (Fig. 2). TSS increased slightly in the lacustrine zone and decreased in the transitional zone, suggesting the event was not of sufficient magnitude to lead to substantive additional erosion in the upstream catchment. Light irradiation increased in both areas (Fig. 4). In the transitional zone, this change was concurrent with a 66% decrease in total phytoplankton biomass and ~50 % decrease in total suspended solid while the lacustrine showed increases in both during this period (Figs. 4 & 6). Though the phytoplankton community composition in the transitional zone did not change, the lacustrine showed a marked increase in the cryptophyte *Katablepharis ovalis* (P6568) for the rest of the sampling period. Total cyanobacterial toxin concentrations declined by ~75% in the transitional zone, while more muted declines were apparent in the lacustrine. All MC variants were replaced with AP-A as the dominant metabolite at all sites by 25 July (DOY 206).

## Discussion

Globally, increases in extreme rainfall events are predicted to outpace changes in total precipitation (Allen and Ingram 2002, IPCC 2007, Soulis et al. 2016). The impacts of extreme rainfall on cyanobacterial blooms are inherently complex (Reichwaldt and Ghadouani 2012, Wood et al. 2017) with the overall effect regulated by parameters including intensity, volume of water inflow, and timing with respect to seasonality (reviewed in Reichwaldt and Ghadouani 2012). The limited studies investigating such extreme rainfall events on cyanobacterial bloom development have identified changes to flushing rates, water column mixing, and nutrient inputs from rainfall events as the main abiotic factors affecting cyanobacteria and phytoplankton communities (Bouvy et al. 2003, Reichwaldt and Ghadouani 2012) – all factors that appeared to impact the bloom dynamics in Conestogo Lake.

The record rainfall event effectively created a new high-P regime as a result of the significant influx of P arriving from the Conestogo River, and near complete replacement of the reservoir volume. Soluble reactive phosphorus concentrations increased ~7-27 fold in the epilimnion following the flooding. Substantial increases in P loading, such as these, are common following heavy rainfall, particularly if they are preceded by warm, dry periods (Jones & Poplawski, 1998). For example, in Australian reservoirs, a 440 mm rainfall in a 3-day period resulted in an input equivalent to 400% of the average annual in-lake P mass (Jones and Poplawski 1998). In our study system, the Upper Conestogo watershed, which is predominantly Tavistock Till, is particularly prone to erosion and serves as a key mechanism for particulate P transport (Loomer & Cooke, 2003; Macrae, English, Schiff, & Stone, 2007). High flow events can also be important to transport of soluble P species, as P is desorbed from particulate forms, or released from soils.

Though surface water temperatures remained similar following the June extreme rainfall event, the bottom sediments warmed from 13 to 18°C. While we cannot isolate the effects of thermal and biogeochemical changes in bloom initiation, we note that nitrogen concentrations changed little as a result of the flooding. This catchment has chronic high N concentrations (3 - 5 mg NO_3_-N L^-1^), associated with long-term intensive agricultural usage within the catchment. In addition to the prevalence and eventual dominance by an N-fixing cyanobacterium, the lack of change in N suggests that it was not likely to be the key factor leading to early bloom initiation.

Instead, it appears that the excessive supply of P initiated the development of the observed weakly toxic early-summer cyanobacterial bloom (Figs 5 and 6). Typically, *Aphanizomenon flos-aquae* dominated blooms do not develop in the Conestogo Reservoir until late summer (S. Cooke, pers communication), ~35 to 40 days after our documented extreme event. Interestingly, the bloom originating in the transitional zone was not only initiated a week earlier than in the lacustrine, but it was also sustained through the typical bloom period, highlighting the potential for elongation of the bloom season and risk associated with extreme rainfall events. However, the intensity of the transitional bloom was directly influenced by the mid-summer rainfall event, which flushed both biomass and toxins. The event was not of sufficient volume to have similar effects on the deeper, larger volume lacustrine zone.

The environmental factors driving toxin production and their seasonal dynamics are quite complex. We detected low quantities of three MC variants, MC-LR, -YR, and -RR in the early-summer cyanobacterial bloom that dissipated following the mid-summer event. Of these, only MC-LR was significantly correlated with the low biomass of a single taxa (*M. aeruginosa*). However, because it difficult to assign strong relationships between the toxins and cyanobacterial species without direct molecular studies (Vezie et al., 1998), it remains possible that there could be the presence of a rare, toxin-producing species or strain, or multiple taxa involved in toxin production.

MC variants are not the only cyanobacterial compounds capable of impacting human health. Several studies using cyanobacterial extracts have reported harmful and/or toxic effects that could not be explained solely by microcystin concentration or presence, suggesting the possibility of other toxic compounds (Keil et al. 2002, Teneva et al. 2005, Baumann and Jüttner 2008, Smutná et al. 2014, Lenz et al. 2019). Although not often reported, improved analytical techniques have identified numerous other bioactive compounds such as cyanopeptolins and anabaenopeptins that are both detectable in freshwaters and often produced simultaneously with microcystin variants (Harada et al. 1995, Welker and Von Döhren 2006, Gkelis et al. 2015, Beversdorf et al. 2017, 2018) at similar or greater levels (Janssen 2019). For example, Beversdorf *et al*. (2017) reported an average of 0.65 μg L^-1^ total microcystin in Lake Koshkonong, Wisconsin, while anabaenopeptin-B and -F were measured at 6.56 μg L^-1^ combined. Though there are no case studies of human toxicity caused by anabaenopeptins, the compound inhibits carboxypeptidase A, and like microcystin, also inhibits protein phosphatases with slightly overlapping inhibitory concentration ranges (Honkanen et al. 1990, Sano et al. 2001, Spoof et al. 2016). The concentrations of these toxins that could affect human health are unknown, which has prevented the development of recreational and drinking water regulations and/or advisories for these potentially toxic compounds. Like some microcystins, the ecological effects of anabaenopeptins may be observed at relatively low concentrations. A recent study by Lenz *et al*. (2019) reported induced toxicity by low concentrations (10 μg L^-1^) of anabaenopeptins, including AP-A, on the nematode *Caenorhabditis elegans* resulting in reduced reproduction, reduced lifespan, and delayed hatching. Though compounds like AP-A have been previously considered non-toxic, they may represent a new class of emerging toxins, whose potential impacts to human health and toxicity to aquatic organisms require immediate attention and therefore, inclusion in risk assessment for lake mitigation and monitoring programs.

While increasing bloom incidence globally has been associated with elevated nutrient loads and with warming (Ho & Michalak, 2019; Paerl & Otten, 2016), more attention is needed to understand the role of extreme events in altering bloom risk, and how these events may differ in their seasonal impacts, and on spatially complex but crucial water resources, such as reservoirs. The environmental factors driving cyanobacterial blooms are inherently complex and dynamic, making predictions about toxicity and impact on end-user health and safety difficult to obtain across diverse ecosystems. Extreme disturbance events, like the early summer event described here, will only make that task more difficult since they are not only predicted to increase but will also likely mask historical drivers of changes in bloom risk. Managers cannot effectively manage a future ecosystem based on averages of the past. Instead, a holistic understanding of multiple drivers of cyanobacterial bloom risk, accounting for both extreme events, and reservoir structure, is required. Within this region, we hypothesize that early season events, have the greatest potential to worsen bloom risk by extending the bloom season as shown here. Later season events can increase annual nutrient loads but may also help flush existing blooms. Here, this appears to occur during the late season runoff event with apparent effects on toxins and cyanobacterial biomass in the transitional zone, with only minimal change apparent in the lacustrine zone.

## Author’s contributions

MLL, HMB, DS, SS, SLS, and JJV conceived of the ideas and designed methodology; MLL, DS, SS collected the data; MLL analyzed the data and lead the writing of the manuscript. All authors contributed critically to the drafts and gave final approval for publication.

## Acknowledgments

Funding for this work was provided by the Canada First Research Excellence Fund program. This was a part of the Global Water Futures Initiative FORMBLOOM: FORecasting tools and Mitigation options for diverse BLOOM-affected lakes as well as the ATRAPP project (Algal Blooms, Treatment, Risk Assessment, Prediction and Prevention through Genomics) from Genome Canada and Genome Québec. We thank S. Cooke, D. McFadden, and the Grand River Conservation Authority for their cooperation and knowledgeable insight, R. Elgood, E. McQuay, T. Cornell, B. Gruber, S. Sine, and M. Soares-Paquin for assistance with sample collection and analysis, J. Atkins for her GIS assistance and map production, and the members of the Venkiteswaran and Schiff lab groups for their reviews of manuscript drafts and friendly discussion.

## Data availability statement

Data and code that support the analysis and findings in this study are available in figshare at http://doi/10.6084/m9.figshare.7811963.

