## Supporting Information for "Extreme rainfall drives early onset cyanobacterial bloom"

**Title:** Extreme rainfall event drives early onset cyanobacterial bloom

Figure S1. Proportional biomass of the six major phytoplankton groups in each sampling location. Rainfall events are marked by the vertical grey bars.

Figure S2: Cyanobacterial proportion in the phytoplankton community in the transitional and lacustrine sites throughout the sampling period.

**Figure S1**

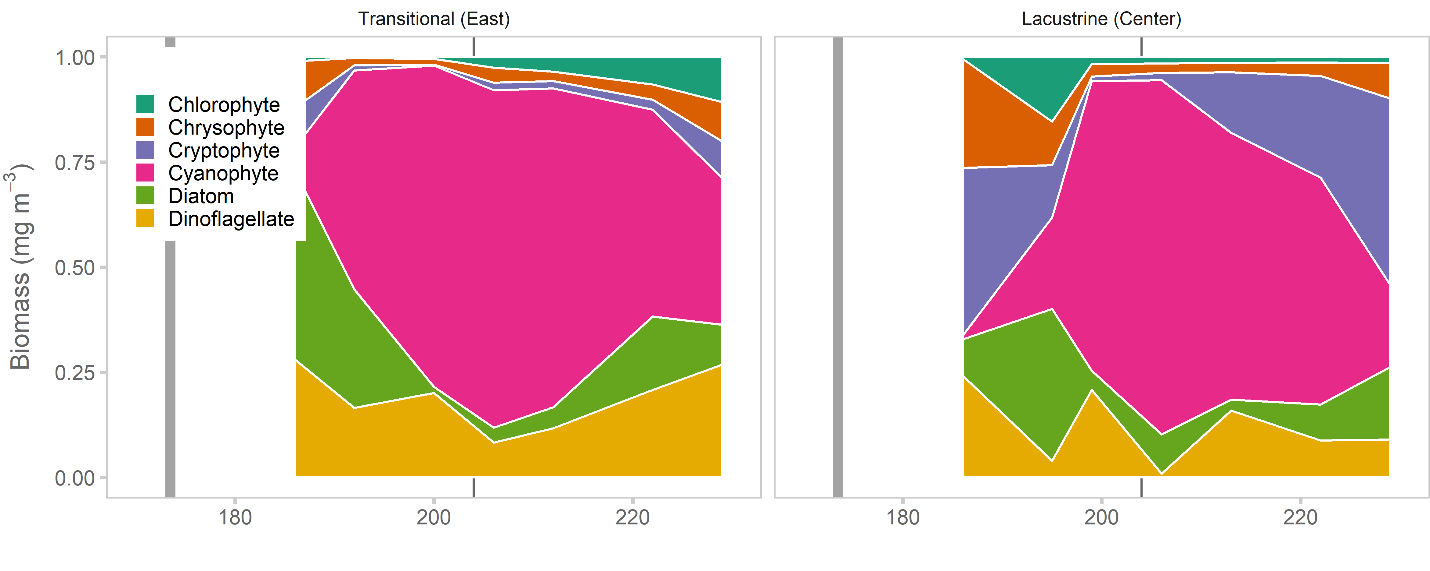

**Figure S2**

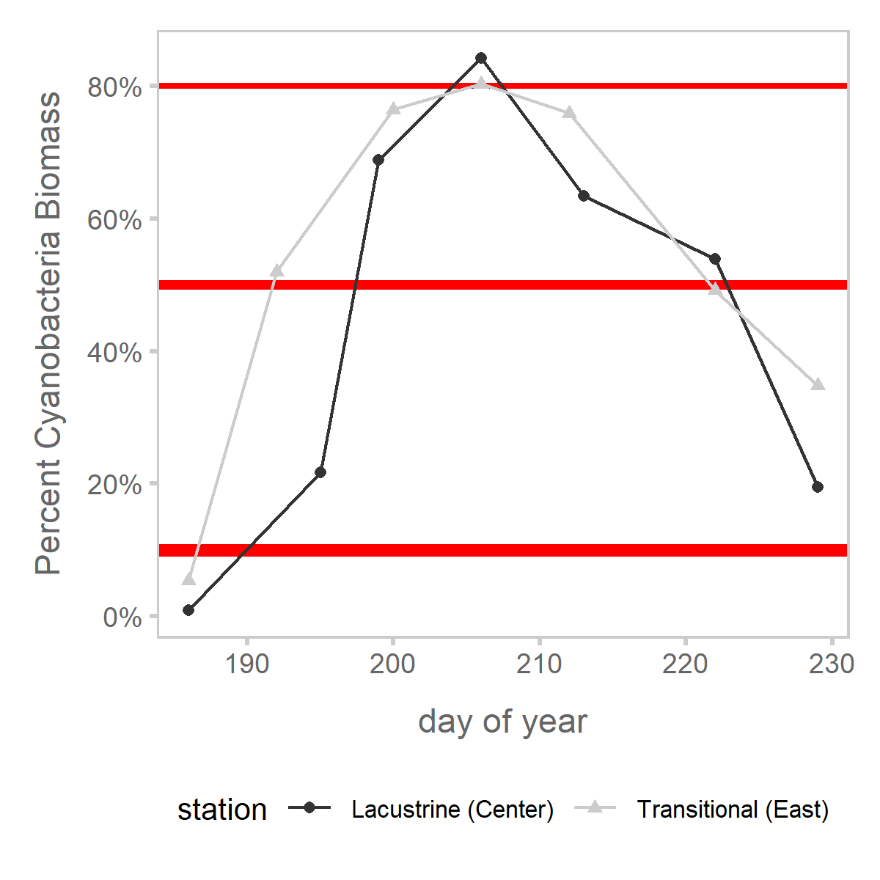

**Table S1** Cyanobacterial metabolite standards used in this study.

| **Compound** | **Abbreviation** | **Manufacturer** | **Catalogue Number** |
| --- | --- | --- | --- |
| Anatoxin-a | ATX | Enzo Life Science | BML-C118-0001 |
| Homoanatoxin-a | HATX | Abraxis | 142926-86-1 |
| Cylindrospermopsin | CYN | Enzo Life Science | ALX-350-149-C025 |
| Microcystin-LR | MC-LR | Enzo Life Science | ALX-350-012-C100 |
| [D-Asp3]Microcystin-LR | [Asp3] MC-LR | Enzo Life Science | ALX-350-173-C025 |
| [Asp3Dha7]-Microcystin-LR | [Asp3Dha7]-MC-LR | NRC | 134842-07-2 |
| Microcystin-RR | MC-RR | Enzo Life Science | ALX-350-043-C050 |
| Microcystin-YR | MC-YR | Enzo Life Science | ALX-350-044-C025 |
| [D-Asp3]Microcystin-RR | [Asp3] MC-RR | Enzo Life Science | ALX-350-168-C025 |
| Microcystin-LA | MC-LA | Enzo Life Science | ALX-350-096-C025 |
| Microcystin-LY | MC-LY | Enzo Life Science | ALX-350-148-C025 |
| Microcystin-LW | MC-LW | Enzo Life Science | ALX-350-080-C025 |
| Microcystin-LF | MC-LF | Enzo Life Science | ALX-350-081-C025 |
| Microcystin-WR | MC-WR | Enzo Life Science | ALX-350-167-C025 |
| Microcystin-HtyR | MC-HtyR | Enzo Life Science | ALX-350-174-C025 |
| Microcystin-HilR | MC-HilR | Enzo Life Science | ALX-350-177-C025 |
| Cyanopeptolin A | CAP | Cyano Biotech GmbH | ALX-350-149-C025 |
| Anabaenopeptin A | AP.A | Enzo Life Science | ALX-350-183-C100 |
